# Colistin resistance in *Escherichia coli* confers protection of the cytoplasmic but not outer membrane from the polymyxin antibiotic

**DOI:** 10.1101/2021.07.21.453182

**Authors:** Madeleine Humphrey, Gerald J. Larrouy-Maumus, R. Christopher D. Furniss, Despoina A. I. Mavridou, Akshay Sabnis, Andrew M. Edwards

**Affiliations:** MRC Centre for Molecular Bacteriology and Infection, Imperial College London, Armstrong Rd., London, SW7 2AZ, UK; Centre for Bacterial Cell Biology, Biosciences Institute, Faculty of Medical Sciences, Newcastle University, Baddiley-Clark Building, Richardson Road, Newcastle upon Tyne, NE2 4AX; Science for Life Laboratory, Department of Molecular Biosciences, The Wenner-Gren Institute, Stockholm University, 106 91 Stockholm, Sweden; Department of Molecular Biosciences, University of Texas at Austin, Austin, 78712, Texas, USA

**Keywords:** Colistin, polymyxin, *E. coli*, lipopolysaccharide, resistance, MCR

## Abstract

Colistin is a polymyxin antibiotic of last resort for the treatment of infections caused by multi-drug resistant Gram-negative bacteria. By targeting lipopolysaccharide (LPS), the antibiotic disrupts both the outer and cytoplasmic membranes, leading to lysis and bacterial death. Colistin resistance in *Escherichia coli* occurs via mutations in the chromosome or the acquisition of mobilised colistin resistance (*mcr*) genes. Both these colistin resistance mechanisms result in chemical modifications to the LPS, with positively charged moieties added at the cytoplasmic membrane before the LPS is transported to the outer membrane. We have previously shown that MCR-1-mediated LPS modification protects the cytoplasmic but not the outer membrane from damage caused by colistin, enabling bacterial survival. However, it remains unclear whether this observation extends to colistin resistance conferred by other *mcr* genes, or resistance due to chromosomal mutations. Using a panel of clinical *E. coli* that had acquired *mcr* -1, -1.5, -2, -3, -3.2 or -5, or had acquired polymyxin resistance independently of *mcr* genes, we found that almost all isolates were susceptible to colistin-mediated permeabilisation of the outer, but not cytoplasmic, membrane. Furthermore, we showed that permeabilisation of the outer membrane of colistin resistant isolates by the polymyxin is in turn sufficient to sensitise bacteria to the antibiotic rifampicin, which normally cannot cross the LPS monolayer. These findings demonstrate that colistin resistance in *E. coli* is typically due to protection of the cytoplasmic but not outer membrane from colistin-mediated damage, regardless of the mechanism of resistance.

## Introduction

The highest priority antibiotic resistant pathogens identified by the World Health Organisation (WHO) are multi-drug-resistant carbapenemase-producing Gram-negative bacteria, including the Enterobacteriaceae, *Pseudomonas aeruginosa and Acinetobacter baumannii* [1]. The Enterobacteriaceae include *Escherichia coli*, which is responsible for over 30,000 cases of bacteraemia in the UK annually [2], while also being the most common causative agent of urinary tract infections [3].

Resistance to first- and second-line antibiotics frequently necessitates the use of drugs of last resort such as the polymyxins, colistin and polymyxin B, which were used to treat 28% of infections caused by carbapenem-resistant Enterobacteriaceae in the USA in the 12 months to January 2019 [4]. Colistin was discovered in 1947 [5] and is only active against Gram-negative bacteria, including most members of the Enterobacteriaceae and other common non-fermentative Gram-negative bacteria, including *P. aeruginosa* and *A. baumannii* [6]. The drug was initially widely prescribed, but its use quickly dwindled due to its lack of efficacy and side effects such as nephrotoxicity and neurotoxicity [7]. Despite these limitations, colistin is considered by the WHO to be a “highest priority critically important antimicrobial for human medicine” because of its ability to treat infections caused by bacteria that are otherwise resistant to antibiotic treatment [8].

The structure of colistin consists of a cationic peptide ring made up of seven amino acids connected to a hydrophobic lipid tail [7,9,10]. The cationic peptide ring of colistin binds the negatively charged lipid A moiety of LPS in the outer membrane (OM), destabilising the cation bridges between LPS molecules and causing the membrane to be disrupted. The acyl tail of colistin is then able to interact with the fatty acid tails of lipid A, which further damages the OM [9]. Colistin then crosses the OM via a process termed ‘self-directed uptake’ to enter the periplasm [9]. Once in the periplasm, colistin likely binds to various macromolecules before it reaches the cytoplasmic membrane (CM) [11]. Subsequently, the antibiotic interacts with LPS in the CM as it is being trafficked to the OM, resulting in CM permeabilisation [12]. It is this interaction with CM LPS that is key to the bactericidal action of colistin, since destabilisation of the CM leads to cell lysis and bacterial death [12,13].

Resistance to colistin can be acquired through chromosomal point mutations, particularly in genes encoding two-component regulatory systems, such as PhoPQ and PmrAB/BasRS [14-17]. These mutations result in the constitutive expression of *eptA* and the *arnBCADTEF* operon, which, in turn, leads to modification of LPS via the addition of 4-amino-4-deoxy-L-arabinose (L-Ara4n) and/or phosphoethanolamine (pEtN) groups to lipid A [17,18,19]. In addition to chromosomal mutations, in 2016, a plasmid-encoded form of colistin resistance was discovered in multiple isolates from both humans and livestock. A single gene, mobilised colistin resistance-1 (*mcr-1*), encoding a pEtN transferase, was shown to confer resistance to polymyxin antibiotics in *E. coli* [20]. Subsequent work demonstrated that *mcr-1* is disseminated globally in a range of different Enterobacteriaceae, particularly *E. coli*, and there are now reports of 10 distinct classes of *mcr*, all of which encode pEtN transferases [6,21,22]. In MCR-mediated colistin resistance, the LPS is solely modified with pEtN, whereas when resistance is conferred by chromosomal mutations, both L-Ara4n and/or pEtN modifications occur [23].

In both MCR-mediated and mutation-mediated colistin resistance, LPS modification occurs in the outer leaflet of the CM [20,24,25]. This results in the presence of modified LPS in both the CM and OM, although not all LPS molecules are modified in either membrane [12,23]. Since both L-Ara4n and pEtN are positively charged, they reduce the anionic charge of lipid A, which is thought to reduce its affinity for the cationic peptide ring of colistin [17,24]. Nonetheless, there is evidence that colistin can still damage the OM of *E. coli* expressing *mcr*-1, most likely due to the presence of unmodified LPS molecules that can be engaged by the polymyxin antibiotic [12,23,26]. By contrast to the damage it caused to the OM, colistin did not permeabilise the CM of an MCR-1-producing strain, explaining the ability of the bacterium to survive polymyxin exposure [12]. This can be explained by the high level of modified LPS in the CM of the MCR-1 strain, along with the low overall abundance of LPS in the CM,

which means there are very few unmodified LPS molecules that colistin can target in the CM [12]. Hence, the MCR-1 pEtN transferase protects the CM, but not OM, of *E. coli* from colistin.

Whilst all MCRs are pEtN transferases located in the CM, it is not known whether observations for MCR-1-expressing bacteria are applicable to bacteria encoding other *mcr* genes. Furthermore, it is not known whether non-MCR-mediated colistin resistance, which commonly results in LPS modified with L-Ara4n and pEtN, similarly confers protection of the CM but not the OM from colistin-mediated damage. This is important to resolve because new approaches aiming to target and overcome colistin resistance may need to be different depending on the mechanism by which resistance is conferred.

## Methods

### Bacterial strains and growth conditions

The *E. coli* strains used in this study are listed in Table 1. Bacterial strains were grown at 37°C with shaking (180 rotations per minute (r.p.m.)) for 18 hours to stationary phase in Luria Broth (LB; Thermo Fisher Scientific, USA). For culture on solid media, bacteria were grown on LB supplemented with 1.5% technical agar (BD Biosciences, USA). Bacterial colony forming units (c.f.u.) were quantified by serial 10-fold dilution of bacterial cultures and plating onto Mueller-Hinton broth (MHB; Sigma-Aldrich, USA) supplemented with 1.5% technical agar. Agar plates were incubated statically in air for 18 hours at 37°C.

**Table 1.** Strains used in this study, the mechanism of colistin resistance, type and degree of LPS modification and susceptibility to colistin. ATCC: American Type Culture Collection.

| *E. coli*<br>Strain | Colistin<br>resistance<br>mechanism | LPS<br>modification | Modified:<br>Unmodified<br>Lipid A Ratio<br>(Whole Cells) | Modified:<br>Unmodified<br>Lipid A Ratio<br>(Spheroplasts) | Colistin<br>MIC ($\mu\text{g mL}^{-1}$) | Source or<br>Reference |
| --- | --- | --- | --- | --- | --- | --- |
| ATCC<br>25922 | None | None | N/A | N/A | 0.5 | ATCC |
| BM16 | None | None | N/A | N/A | 1 | [29] |
| DIN | None | None | N/A | N/A | 1 | [29] |
| CNR 1745 | MCR-1 | pEtN | $2.18 \pm 0.75$ | $3.25 \pm 1.00$ | 2 | [29] |
| CNR<br>20140385 | MCR-1 | pEtN | $1.92 \pm 0.04$ | $2.40 \pm 2.02$ | 2 | [29] |
| CNR 1790 | MCR-1 | pEtN | $1.28 \pm 0.23$ | $1.56 \pm 0.71$ | 2 | [29] |
| 1078733 | MCR-1 | pEtN | $2.59 \pm 1.82$ | $2.28 \pm 0.04$ | 2 | [30] |
| 1256822 | MCR-1.5 | pEtN | $2.80 \pm 0.04$ | $1.82 \pm 0.39$ | 2 | [30] |
| R11 | MCR-2 | pEtN | $1.27 \pm 0.12$ | $1.32 \pm 0.27$ | 2 | [30] |
| 1488949 | MCR-3 | pEtN | $1.94 \pm 0.10$ | $2.02 \pm 0.49$ | 2 | [30] |
| 1267171 | MCR-3 | pEtN | $2.76 \pm 0.35$ | $2.20 \pm 1.20$ | 2 | [30] |
| 1266877 | MCR-3.2 | pEtN | $2.67 \pm 1.81$ | $1.94 \pm 0.31$ | 2 | [30] |
| 1144230 | MCR-5 | pEtN | $1.60 \pm 1.09$ | $1.98 \pm 0.82$ | 2 | [30] |
| CNR 1728 | Chromosomal<br>PmrB (G160E) | pEtN & L-Ara4n | $5.41 \pm 3.10$ | $3.33 \pm 0.23$ | 2 | [29] |
| 1195290 | Chromosomal | pEtN & L-Ara4n | $3.00 \pm 0.00$ | $4.14 \pm 0.65$ | 4 | [30] |
| 1272408 | Chromosomal | pEtN & L-Ara4n | $5.66 \pm 1.35$ | $2.73 \pm 1.11$ | 4 | [30] |
| 1262287 | Chromosomal | pEtN & L-Ara4n | $7.94 \pm 3.31$ | $3.81 \pm 0.40$ | 4 | [30] |
| 1252394 | Chromosomal | pEtN & L-Ara4n | $1.55 \pm 0.31$ | $2.48 \pm 0.16$ | 4 | [30] |
| 1150735 | Chromosomal | pEtN & L-Ara4n | $4.67 \pm 0.71$ | $2.81 \pm 0.68$ | 4 | [30] |

### Determination of minimum inhibitory concentrations of antibiotics

Broth microdilution was used to determine the minimum inhibitory concentration (MIC) of colistin sulphate (Sigma-Aldrich, USA) for each bacterial strain [27]. A range of antibiotic concentrations was prepared by two-fold serial dilution of colistin in 200 µL MHB in a microtitre plate. Stationary phase bacteria diluted 1000-fold in fresh MHB were inoculated into each well of the microtitre plate to achieve a final concentration of 5 × 10^5^ c.f.u. mL^-1^. The MIC was defined as the lowest concentration of antibiotic in which there was no visible growth of bacteria after 18 hours static incubation in air at 37°C. Visible growth was verified by measuring optical density at 595 nm (OD595nm) on a Bio-Rad iMark microplate absorbance reader (Bio-Rad Laboratories, USA). Subsequent MIC assays for rifampicin (Molekula, UK) were run in the absence or presence of colistin at 1 µg mL^-1^ to determine the impact of the polymyxin on rifampicin susceptibility of the colistin resistant clinical isolates.

### OM disruption assay

Disruption of the OM of bacteria was detected using the *N*-phenyl-1-napthylamine (NPN) uptake assay [12]. Bacteria grown to stationary phase overnight were washed three times in MHB by centrifugation (12,300 x *g*, 3 minutes) and resuspension. Washed bacteria were diluted to an OD600nm of 0.5 in 5 mM HEPES buffer (Sigma-Aldrich, USA) and added to a black-walled microtitre plate. NPN (Acros Organics, USA) was diluted in HEPES buffer and added to the relevant wells to achieve a final concentration of 10 μM. Colistin was diluted in HEPES buffer and added to the relevant wells to achieve a final concentration of 4 μg mL^-1^. Fluorescence was measured immediately using a Tecan Infinite M200 Pro microplate reader (Tecan Group Ltd., Switzerland) using an excitation wavelength of 355 nm and an emission wavelength of 405 nm. Measurements were obtained every 30 seconds for 1 hour and averaged to give mean fluorescence. The degree of OM permeabilisation was calculated as the NPN uptake factor [28]:

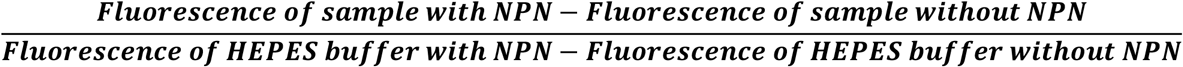

### CM disruption assay

Stationary phase bacteria grown overnight were washed in MHB, as described above for the OM disruption assay, and added at a final inoculum of 10^8^ c.f.u. mL^-1^ to 3 mL MHB containing 4 µg mL^-1^ colistin. These cultures were incubated at 37°C with shaking (180 r.p.m.) for 1 hour before aliquots (200 µL) were taken, bacteria were isolated by centrifugation (12,300 x *g*, 3 minutes) and resuspended in 200 µL phosphate-buffered saline (PBS; Sigma-Aldrich, USA). Resuspended cells (200 µL) were added to a black-walled microtitre plate and propidium iodide (PI; Sigma-Aldrich, USA) was added to achieve a final concentration of 2.5 µM. After 3 hours incubation with PI at room temperature, relative fluorescence units (r.f.u.) were determined using a Tecan Infinite M200 Pro microplate reader with an excitation wavelength of 535 nm and an emission wavelength of 617 nm. Fluorescence was blanked against MHB alone, and differences in fluorescence due to variation in cell number caused by the growth inhibitory effects of colistin were corrected for cell number using OD600nm readings.

### Determination of bactericidal activity of colistin

Stationary phase bacteria were washed in MHB as described above and added to 3 mL MHB containing 4 µg mL^-1^ colistin to give an inoculum of 10^8^ c.f.u. mL^-1^. These cultures were incubated at 37°C with shaking (180 r.p.m.) for 8 hours, with aliquots (200 µL) taken at 0, 2 and 8 hours and survival determined by serial dilution in PBS followed by enumeration of c.f.u. on Mueller-Hinton agar (MHA). Percentage survival was calculated relative to the starting inoculum.

### Determination of bacterial lysis

Stationary phase bacteria were washed with PBS and added at a final inoculum of 10^8^ c.f.u. mL^-1^ to 3 mL MHB containing 4 µg mL^-1^ colistin. Cultures were incubated at 37°C with shaking (180 r.p.m.) for 8 hours. Aliquots (200 µL) were taken at 0, 2 and 8 hours and added to a microtitre plate, with OD595nm measurements subsequently obtained on a Bio-Rad iMark microplate absorbance reader. Fold change in OD595nm readings was calculated relative to the 0 hours measurements.

### LPS characterisation by MALDIxin assay

LPS modifications were detected and quantified using MALDI-TOF mass spectrometry-based lipidomics as described previously [12,23]. Mild-acid hydrolysis was performed on 100 μL suspensions of whole bacterial cells or spheroplasts by adding 100 μL of acetic acid (2 % v/v) and incubating the mixture at 98°C for 30 minutes. Hydrolysed cells/spheroplasts were centrifuged at 17,000 *x g* for 2 minutes, the supernatant was discarded, and the pellet was washed 3 times with 300 μL of ultrapure water. A volume of 0.4 μL of this suspension was loaded onto the MALDI target plate and immediately overlaid with 1.2 μL of Norharmane matrix (Sigma-Aldrich) solubilized in chloroform/methanol (90:10 v/v) to a final concentration of 10 mg mL^-1^. For external calibration, 0.5 µL of calibration peptide was loaded along with 0.5 µL of the given calibration matrix (peptide calibration standard II, Bruker Daltonik, Germany). The samples were loaded onto a MSP 96 target polished steel BC (Bruker Part-No. 8280800).

The bacterial suspension and matrix were mixed directly on the target by pipetting and the mix dried gently under a stream of air. The spectra were recorded in the linear negative-ion mode (laser intensity 95%, ion source 1 = 10.00 kV, ion source 2 = 8.98 kV, lens = 3.00 kV, detector voltage = 2652 V, pulsed ion extraction = 150 ns). Each spectrum corresponded to ion accumulation of 5,000 laser shots randomly distributed on the spot. The spectra obtained were processed with default parameters using FlexAnalysis v.3.4 software (Bruker Daltonik, Germany).

The negative mass spectrum was scanned between *m*/*z* 1,100 and *m*/*z* 2,500 in the negative linear ion mode. Manual peak picking at masses relevant to colistin resistance was performed on the obtained mass spectra and the corresponding signal intensities at these defined masses was determined. The percentage of modified lipid A was calculated by dividing the sum of the intensities of the lipid A peaks attributed to addition of L-Ara4n and/or pETN by the intensity of the peaks corresponding to native lipid A.

## Results

### Colistin resistance due to chromosomal mechanisms is associated with a higher proportion of LPS modification compared to acquisition of *mcr* genes

To understand whether colistin resistance in *E. coli* always results in protection of the CM but not the OM, we first assembled a panel of previously described clinical *E. coli* isolates which were resistant to colistin via either acquisition of an *mcr* gene, or via an *mcr*-independent mechanism consistent with chromosomal mutations, as determined by the presence of LPS modified with both L-Ara4N and pEtN (Table 1) [23,29,30]. As expected, lipidomic analysis demonstrated that *E. coli* strains that harboured an *mcr* gene had pEtN-modified LPS [23]. By contrast, colistin-resistant strains lacking *mcr* genes had LPS that was modified by both pEtN and L-Ara4n, representative of resistance that arises due to chromosomal mutations (Table 1) [23]. Three colistin susceptible strains were included as controls and these did not have detectable LPS modifications. Strains positive for colistin resistance mechanisms had colistin minimum inhibitory concentration (MIC) values of 2-4 µg mL^-1^, which were greater than susceptible strains and largely in agreement with previous reports [23] (Table 1). Notably, 5 of the 6 strains resistant to colistin via chromosomal mutations had higher MIC values than *E. coli* strains harbouring *mcr* genes (Table 1).

To understand the basis of this difference in MIC values between *E. coli* isolates carrying *mcr* genes versus chromosomal mechanisms of colistin resistance, we determined the ratio of modified:unmodified LPS of both whole cells and spheroplasts in our strain panel (Table 1). Since most LPS is in the OM [12,24,25], values for whole cells are largely representative of this membrane, while values for spheroplasts are indicative of LPS modification in the CM. Comparison of values revealed significantly greater levels of LPS modification in both the OM and CM for strains resistant due to chromosomal mechanisms relative to *E. coli mcr* strains (Fig. 1). This suggests that there is a relationship between colistin MIC and the proportion of modified LPS, although it was not clear whether this was due to the proportion of modified LPS in the OM, the CM, or both.

**Figure 1.**
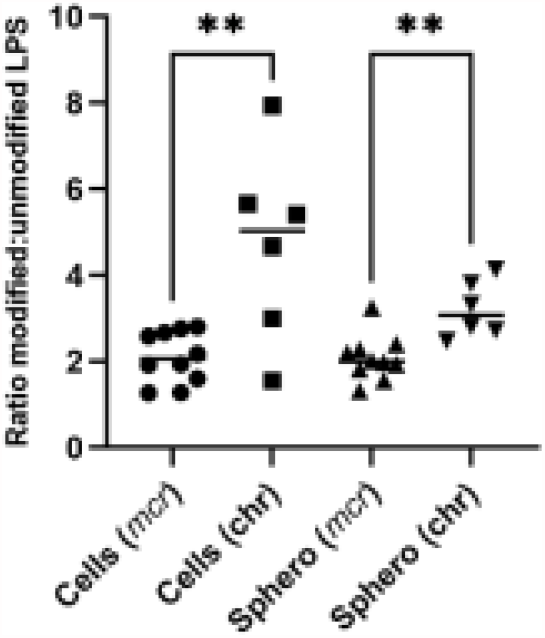
Resistance mediated via chromosomal mutations is associated with higher proportions of modified LPS compared to resistance due to *mcr* gene acquisition. Ratios of modified:unmodified LPS in whole cells (Cells) or spheroplasts (Sphero) from *E. coli* resistant to colistin via *mcr* genes (*mcr*) or via chromosomal mechanisms (chr). Ratios were significantly higher for resistance due to chromosomal mechanisms compared to *mcr* genes for both whole cells and spheroplasts, as determined by a Kruskal-Wallis test, corrected for multiple comparisons via a two-stage linear step-up procedure of Benjamini, Krieger and Yekutieli (**p<0.01).

### Colistin resistance is associated with protection of the CM but not the OM from permeabilisation by the polymyxin antibiotic, regardless of the mechanism of resistance

To test whether colistin permeabilised the OM of our panel of isolates we used the well-established NPN uptake assay [12,26]. The NPN dye becomes fluorescent when bound to phospholipids exposed in strains where the outer LPS monolayer of the OM has been disrupted [28]. Each strain was exposed to colistin at a concentration of 4 µg mL^-1^, the peak serum concentration that can be achieved therapeutically [31], before NPN-mediated fluorescence was measured. This revealed that colistin increased OM permeability throughout the panel of clinical isolates, both in susceptible and colistin-resistant *E. coli* strains, with all but one strain (1078733) exhibiting a significant increase (p<0.05) in NPN uptake in the presence of the polymyxin antibiotic compared to untreated conditions (Fig. 2a). More specifically, the NPN uptake factor of most strains more than doubled in the presence of colistin, and the extent of permeabilisation for many resistant strains was as great as that of the susceptible strains (Fig. 2a).

**Figure 2.**
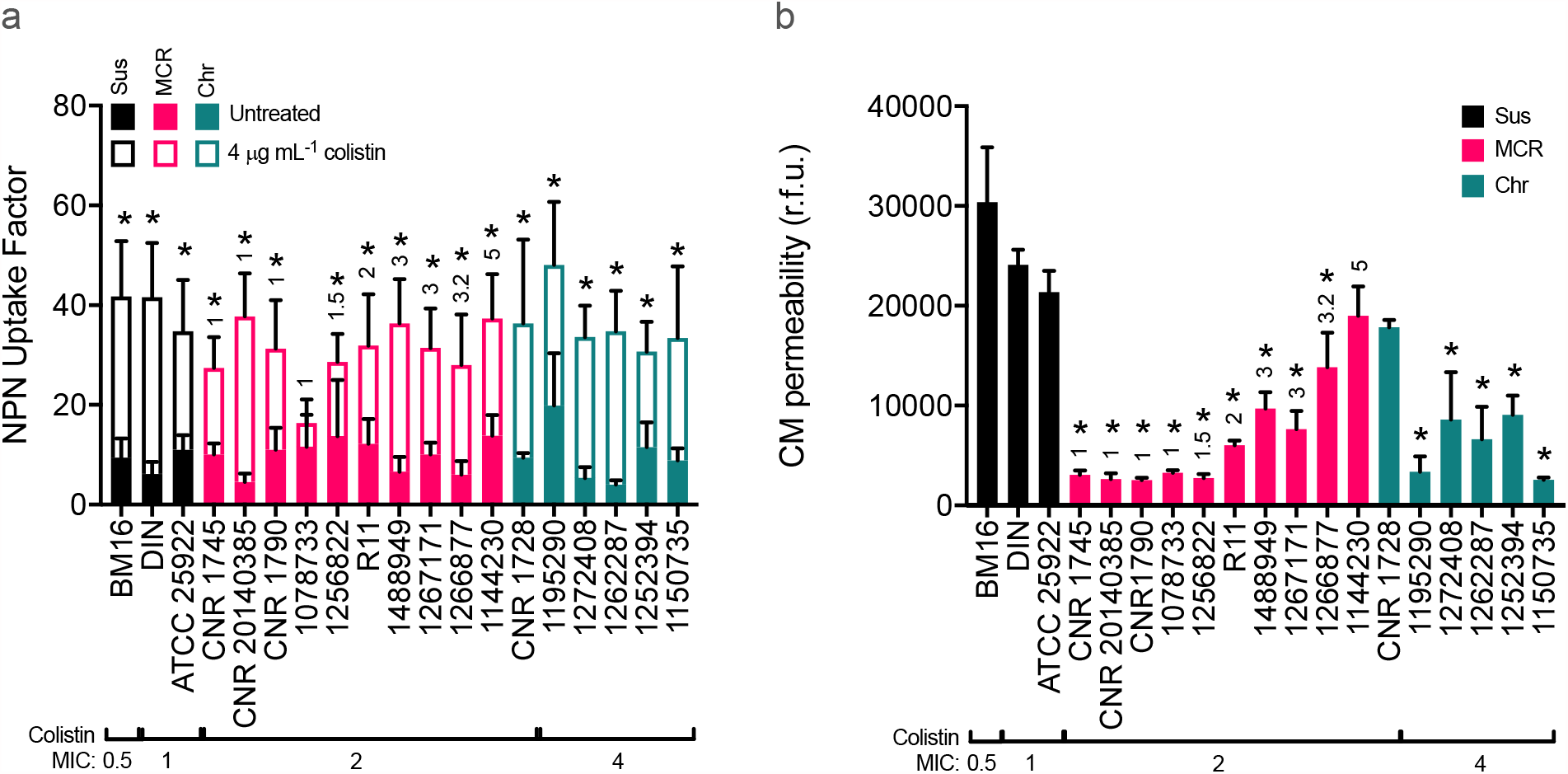
The outer membrane of resistant bacteria is permeabilised by colistin. **a.** Disruption of the outer membrane of *E. coli* clinical isolates with (empty bars) and without (filled bars) incubation with 4 µg mL^-1^ colistin, determined by the uptake of N-phenyl-1-napthylamine (NPN; 10 µM) into *E. coli* cells that were colistin susceptible (Sus), colistin resistant due to the acquisition of *mcr* genes (MCR) or resistant via chromosomal mechanisms (Chr) (n=3, analysed using unpaired multiple t-tests and corrected with Holm-Sidak’s method, *p<0.05 between untreated and 4 µg mL^-1^ colistin for each strain). **b.** Disruption of the cytoplasmic membrane (CM) of clinical isolates after 1 hour incubation with 4 µg mL^-1^ colistin, measured using propidium iodide (2.5 µM) and expressed as relative fluorescence units (r.f.u.) (n=3, analysed by a one-way ANOVA with Dunnett’s post-hoc test, *p<0.05 decrease compared to ATCC 25922). The MCR type is annotated above the *mcr*-harbouring strains.

Having shown that colistin permeabilised the OM of both susceptible and resistant isolates, the next step was to determine the effect of the polymyxin antibiotic on the CM. This was assessed using the membrane impermeant dye PI, which fluoresces when bound to DNA that becomes accessible only when both the OM and CM of the bacteria have been compromised. The colistin-susceptible strains all had a high level of fluorescence from PI in the presence of 4 µg mL^-1^ colistin, showing a large degree of polymyxin-mediated CM permeabilisation (Fig. 2b). By contrast, as shown previously, *mcr*-1-harbouring strains had very little CM disruption in the presence of 4 µg mL^-1^ colistin [12,26]. Bacteria expressing other MCR variants also had significantly less (p<0.05) CM disruption than the EUCAST quality control strain ATCC 25922, except for the *mcr*-5-harbouring strain (1144230), which had a comparable level of PI uptake as the susceptible ATCC 25922 strain. The majority of non-*mcr*-harbouring colistin resistant strains also had significantly lower levels of CM disruption compared to the control strain ATCC 25922, with only one strain (CNR 1728) exhibiting significant CM disruption due to colistin exposure.

Taken together, these data demonstrated that 15 out of 16 colistin resistant strains (94%) experience significant OM but not CM permeabilisation in the presence of a clinically relevant concentration of colistin, supporting the hypothesis that colistin resistance is predominantly due to protection of the CM rather than the OM from the polymyxin antibiotic.

### Colistin-mediated permeabilisation of the OM does not affect bacterial viability

Next, we wanted to understand the consequences of colistin-mediated OM and CM damage for bacterial viability. To do this we exposed the bacteria (∼10^8^ c.f.u. mL^-1^) to 4 µg mL^-1^ colistin and measured survival via c.f.u. counts after 2 and 8 hours. As expected, there was a >1000-fold reduction in c.f.u. counts of the susceptible strains after 2 hours, which was maintained at 8 hours (Fig. 3a,b). By contrast, most of the colistin resistant strains were unaffected by the presence of colistin over 2 hours, although three strains (1266877, 1144230 and CNR 1728) had reduced c.f.u. counts (Fig. 3a). By 8 hours, all but one of the colistin-resistant strains (1144230) had increased c.f.u. counts relative to the start of the assay, indicative of replication in the presence of the antibiotic (Fig. 3b). During these assays we also used OD595nm measurements to detect lysis or growth. As expected for the colistin susceptible strains, all three exhibited reduced OD595nm values at both 2 and 8 hours, indicative of cell lysis (Fig. 3e,f). By contrast, all but 2 of the colistin resistant strains (1266877, 1144230) had increased OD595nm readings after 2 hours (Fig. 3e). By 8 hours, all but one of the resistant strains had >3-fold increase in OD595nm values, relative to the start of the assay, in keeping with increased c.f.u. counts (Fig. 3f). The one exception was strain 1144230, which had only a very small increase in OD595nm readings, and a slight reduction in c.f.u. counts after 8 h exposure to colistin. Notably, this strain also suffered the highest degree of colistin-induced CM damage, further supporting the link between damage to this membrane and lysis/bactericidal activity.

**Figure 3.**
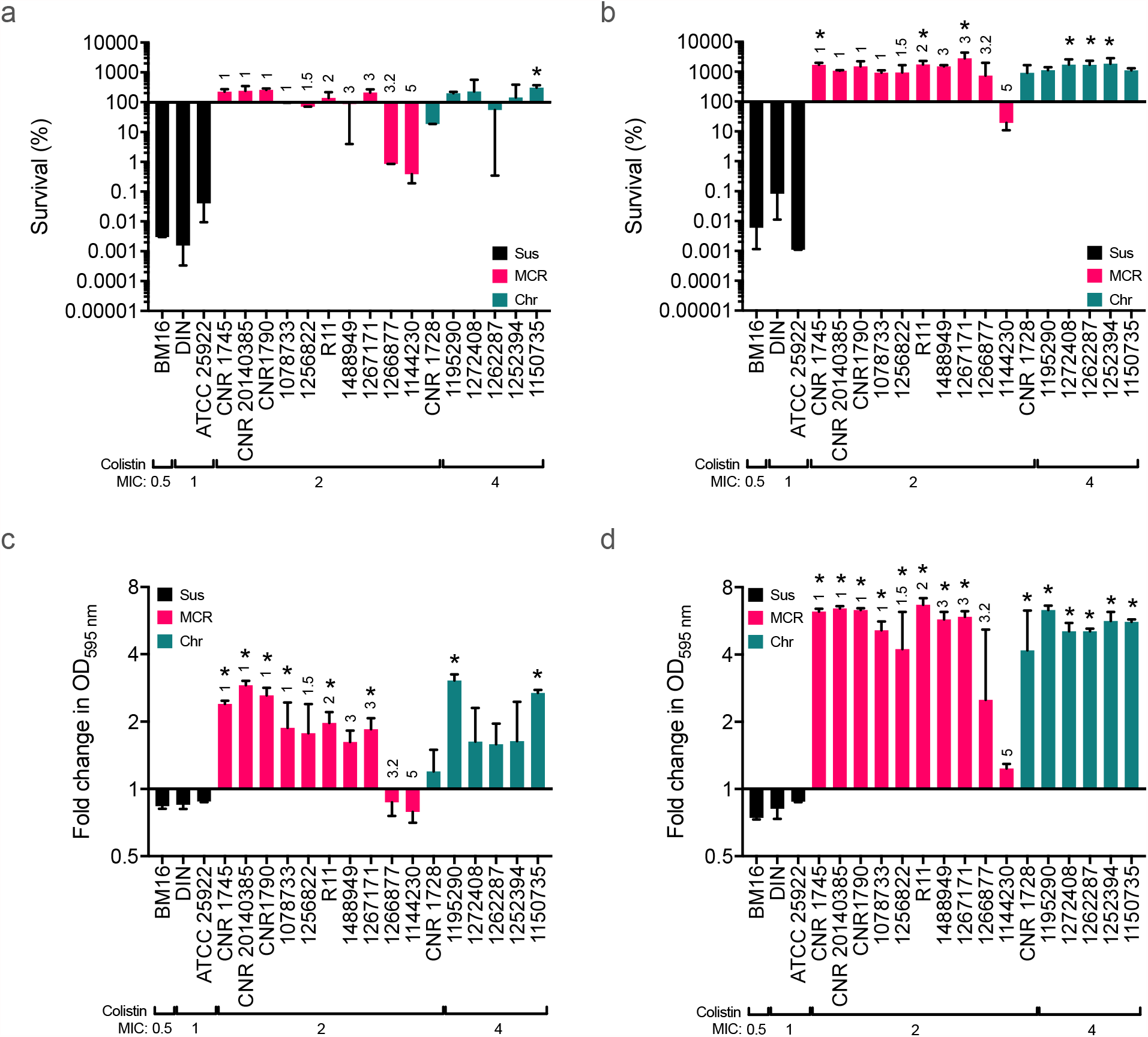
Resistant clinical isolates survive and grow in the presence of colistin. **a, b.** Survival of *E. coli* clinical isolates incubated with 4 µg mL^-1^ colistin at 2 **(a)** and 8 **(b)** hours in *E. coli* that were colistin susceptible (Sus), colistin resistant due to the acquisition of *mcr* genes (MCR) or resistant via chromosomal mechanisms (Chr) (n=3, analysed by a one-way ANOVA with Dunnett’s post-hoc test, *p<0.05 increase compared to ATCC 25922). **c, d.** Lysis or growth of *E. coli* clinical isolates grown with 4 µg mL^-1^ colistin at 2 **(c)** and 8 **(d)** hours, as determined by fold change in OD595nm readings (n=3, analysed by a one-way ANOVA with Dunnett’s post-hoc test, *p<0.05 increase compared to ATCC 25922). The MCR type is annotated above the *mcr*-harbouring strains.

Furthermore, we explored the link between susceptibility of strains to the bactericidal activity of colistin and the degree of CM and OM damage. This revealed a clear correlation between bacterial survival after 2 hours of colistin exposure and the degree of CM, but not OM permeabilisation (Fig. 4a,b). Taken together, these data demonstrate that colistin-mediated permeabilisation of the OM of resistant isolates does not inhibit bacterial replication and provides support for previous work showing that damage to the CM is critical for the bactericidal activity of the antibiotic [12,13].

**Figure 4.**
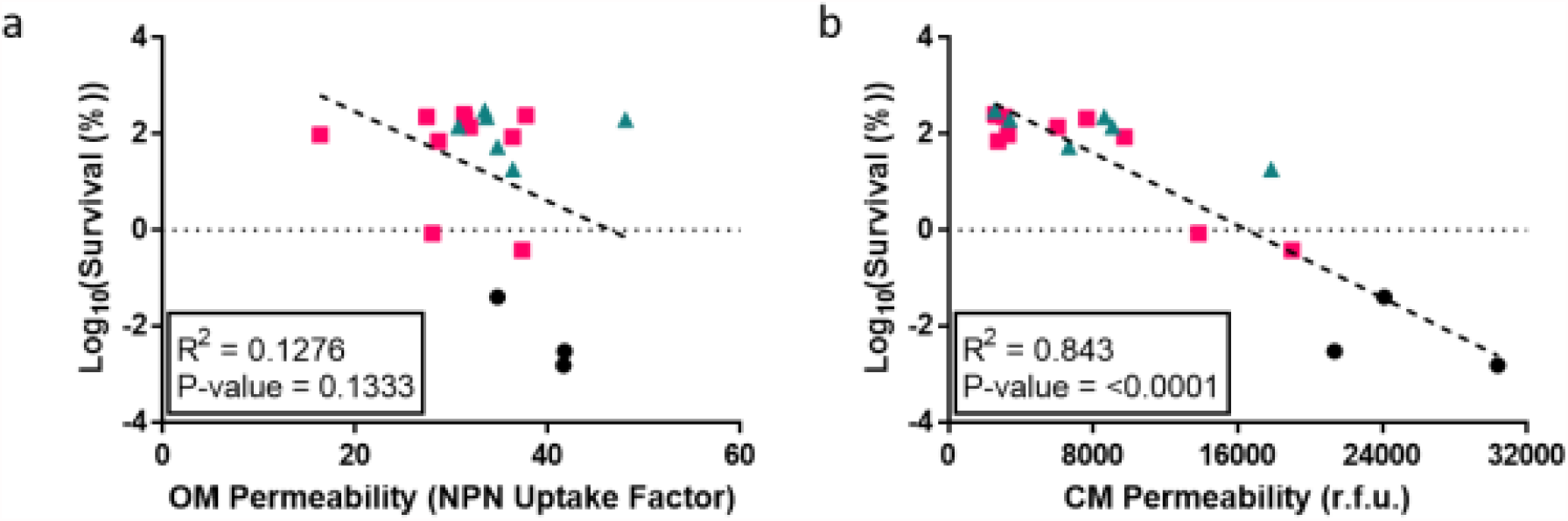
Susceptibility to the bactericidal activity of colistin correlates with CM, but not OM, damage in both resistant and susceptible strains. **a, b.** Scatter plots of bacterial survival after 2 hours exposure to colistin (4 µg mL^-1^) versus OM permeability **(a)** as defined by the NPN uptake factor, or CM permeability **(b)** as measured by fluorescence from propidium iodide binding to chromosomal DNA. Symbols represent the values for *E. coli* strains that were colistin-susceptible strains (Sus, black circles), or colistin resistant due to the acquisition of *mcr* genes (MCR, pink squares) or resistant via chromosomal mechanisms (Chr, blue triangles). Correlations between membrane permeability and survival were assessed by determining the p-value and R^2^ value following linear regression analyses.

### Colistin-mediated OM disruption sensitises polymyxin-resistant bacteria to rifampicin, regardless of the mechanism of resistance

To better understand the nature of the OM permeabilisation caused by colistin and the potential impact of the different resistance mechanisms, we investigated whether polymyxin-mediated OM damage was sufficient to enable the ingress of small molecules that are usually unable to penetrate the LPS monolayer into colistin-resistant bacteria [26]. This was tested using rifampicin, an antibiotic that is not conventionally used to treat *E. coli* infections because of its poor penetration into Gram-negative bacteria. Rifampicin MICs were determined alone and in the presence of 1 µg mL^-1^ colistin, which was sub-inhibitory to the colistin-resistant isolates in our panel (Table 1). Since this concentration of the antibiotic was inhibitory to the growth of the susceptible strains, these were omitted from the analysis (Fig. 5). The use of a sub-inhibitory concentration of colistin ensured that any growth inhibition was due only to the effects of rifampicin. The MICs of rifampicin alone against the panel of *E. coli* isolates ranged from 4 to 32 µg mL^-1^, with the majority falling at 16 µg mL^-1^ (Fig. 5). However, when 1 µg mL^-1^ colistin was added to the assay, the rifampicin MIC was reduced by at least 4-fold (p<0.05) for all but one strain (1488949) (Fig. 5). In particular, for strains 1144230, CNR 1728, 1252394 and 1150735, colistin-mediated OM disruption reduced the rifampicin MIC by 128-fold (Fig. 5). This demonstrates that whilst colistin-mediated permeabilisation of the OM does not have a detrimental effect on bacterial growth, it is sufficient to enable ingress of a small molecule antibiotic. Interestingly, some of the largest decreases in rifampicin MIC were observed for strains with the highest colistin MICs.

**Figure 5.**
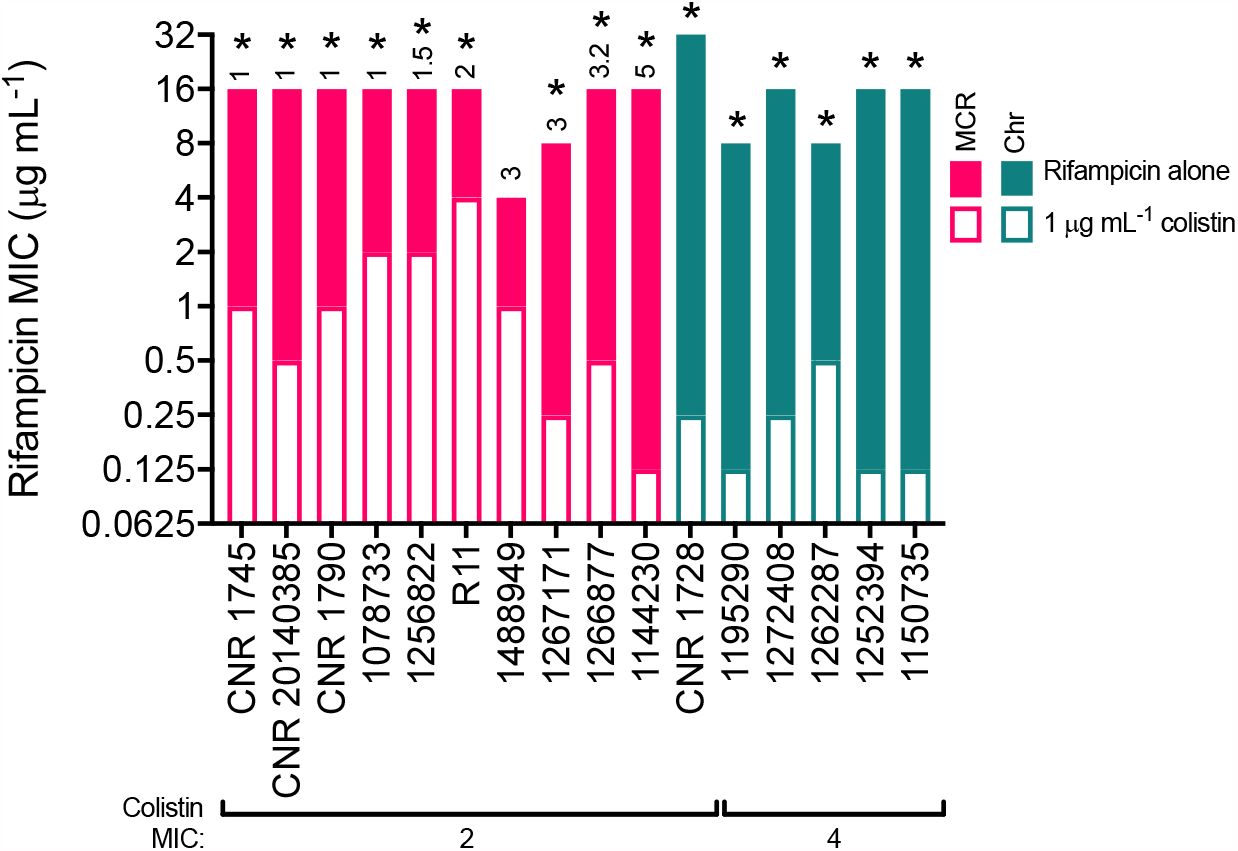
Colistin resistant strains become susceptible to rifampicin when the polymyxin is present at a sub-inhibitory concentration. The minimum inhibitory concentration (MIC) of rifampicin required to inhibit growth of *E. coli* that were resistant to colistin due to the acquisition of *mcr* genes (MCR) or via chromosomal mechanisms (Chr) when incubated in Mueller-Hinton broth (MHB) alone (filled bars) or in MHB containing a sub-inhibitory concentration of colistin (1 µg mL^-1^, empty bars) (n=3, data are presented as the median value, analysed using unpaired multiple t-tests and corrected with Holm-Sidak’s method, *p<0.05 between rifampicin alone and rifampicin with 1 µg mL^-1^ colistin for each strain). The MCR type is annotated above the *mcr*-harbouring strains.

These findings provide additional evidence that colistin resistance is associated with protection of the CM, and not the OM, from the polymyxin antibiotic and are in keeping with recent work showing synergy of colistin with rifampicin against colistin-resistant *E. coli* and *Klebsiella* strains [26,32].

## Discussion

We have shown recently that colistin damages the CM of bacteria by targeting LPS as it is trafficked to the OM, leading to bacterial death and lysis [12]. In addition, we found that MCR-1-mediated colistin resistance protects *E. coli* against damage to the CM via modification of LPS with pEtN [12], but this does not prevent OM permeabilisation. However, it was not known if these findings could be generalised to bacteria with other mechanisms of colistin resistance, such as *mcr* genes belonging to different families or resistance due to chromosomal mutations [12].

The data presented here provide strong evidence that colistin resistance acquired by different mechanisms is associated with protection of the CM, but not the OM, from permeabilisation caused by the polymyxin antibiotic. It is important to note that despite all MCRs being pEtN transferases [21], there were noticeable differences between the ability of the different *mcr* gene families in facilitating *E. coli* survival and growth in the presence of colistin. MCR-1 was consistently shown to be the most effective MCR at conferring protection from colistin, as shown by the low degree of CM permeabilisation by colistin for these strains. This enabled the highest levels of survival and growth of these *E. coli* cells during colistin exposure. By contrast, MCR families 2-5 were less protective against colistin, with higher degrees of CM permeability and reduced survival and growth in the presence of colistin compared to *mcr*-1-harbouring strains. This was especially noticeable for the *mcr*-5-expressing strain (1144230), which had a similar degree of colistin-mediated CM damage to that of the susceptible control strain ATCC 25922. This high level of CM permeability caused a 2-log decrease in survival of strain 1144230 after 2 hours of colistin exposure, and by 8 hours the bacteria had still not recovered back to the starting inoculum. It is not clear why MCR-1 confers greater protection from colistin than other MCRs, but many factors may be involved, including protein production levels, codon usage, promoter strength and protein stability. However, the fact that *mcr-*1 conferred the most protection against colistin out of the MCRs tested in this study may provide an explanation for why *mcr*-1 is the most widely disseminated plasmid-mediated colistin resistance determinant [22].

The finding that colistin permeabilises the OM of all colistin resistant *E. coli* strains may be clinically significant, as it provides a route by which these bacteria can be sensitised to antibiotics that would otherwise be ineffective. Whilst new antibiotics are becoming available to combat carbapenemase-producing Enterobacteriaceae in high-income countries, these are unlikely to be available in low- and middle-income countries. Therefore, the emergence of multi-drug-resistant bacteria necessitates the development of new approaches that employ existing and widely available antibiotics.

Our findings are in keeping with previous reports [26,32] that show that colistin synergises with rifampicin, and suggest that these two antibiotics may provide a useful combination therapy approach that would be cheap and available in both high- and low-income countries. Furthermore, the addition of azithromycin to produce a triple-drug combination has been shown to provide even more efficient synergy against an *mcr*-1-harbouring *E. coli* [33]. However, there is very little clinical data assessing the efficacy of colistin and rifampicin in combination. A trial comparing treatment of extensively drug resistant *A. baumannii* with colistin alone or in combination with rifampicin, found that the combination was more efficacious in eradicating infection, although this did not reduce the overall 30-day mortality rate [34]. Whilst the lack of a significant reduction in death is disappointing, participants in the study population were extremely ill, typically with severe co-morbidities. It may be that earlier introduction of combination therapy may be beneficial at reducing the mortality rate in these patients, and there may also be value in using colistin and rifampicin against more acute infections such as bacteraemia.

In summary, the data described here support previous work based on MCR-1-expressing *E. coli*, by showing that colistin permeabilises the OM, but usually not the CM, of *E. coli* strains that are resistant to colistin regardless of whether resistance is due to MCR or non-MCR-mediated mechanisms.

## Acknowledgements

Dr Laurent Dortet is gratefully acknowledged for the kind gift of the BM16, DIN, CNR 1745, CNR 20140385, CNR 1790, R11 and CNR 1728 isolates. IHMA Inc. Schaumburg are thanked for the kind gift of the *E. coli* 1144230 isolate. A.S. is supported by a PhD studentship funded by a Medical Research Council Doctoral Training Award to Imperial College London (MR/N014103/1). D.A.I.M. and R.C.D.F. gratefully acknowledge funding from an MRC Career Development Award (MR/M009505/1). R.C.D.F. is supported by a Fellowship from the Wenner-Gren Foundations (UPD2019-0174). G.J.L-M. is funded by the MRC Confidence in Concept Fund and a ISSF Wellcome Trust Grant (105603/Z/14/Z). A.M.E. acknowledges support from the National Institute for Health Research (NIHR) Imperial Biomedical Research Centre (BRC).

